# WTAP targets the METTL3 m^6^A-methyltransferase complex to cytoplasmic hepatitis C virus RNA to regulate infection

**DOI:** 10.1101/2022.06.27.497872

**Authors:** Matthew T. Sacco, Katherine M. Bland, Stacy M. Horner

## Abstract

Modification of the hepatitis C virus (HCV) positive-strand RNA genome by N6-methyladenosine (m^6^A) regulates the viral lifecycle. This lifecycle takes place solely in the cytoplasm, while m^6^A addition on cellular mRNA takes place in the nucleus. Thus, the mechanisms by which m^6^A is deposited on the viral RNA have been unclear. In this work, we find that m^6^A modification of HCV RNA by the m^6^A-methyltransferase proteins METTL3 and METTL14 is regulated by WTAP. WTAP, a predominantly nuclear protein, is an essential member of the cellular mRNA m^6^A-methyltransferase complex and known to target METTL3 to mRNA. We found that HCV infection induces localization of WTAP to the cytoplasm. Importantly, we found that WTAP is required for both METTL3 interaction with HCV RNA and for m^6^A modification across the viral RNA genome. Further, we found that WTAP, like METTL3 and METTL14, negatively regulates the production of infectious HCV virions, a process that we have previously shown is regulated by m^6^A. Excitingly, WTAP regulation of both HCV RNA m^6^A modification and virion production were independent of its ability to localize to the nucleus. Together, these results reveal that WTAP is critical for HCV RNA m^6^A modification by METTL3 and METTL14 in the cytoplasm.

**IMPORTANCE:** Positive-strand RNA viruses such as HCV represent a significant global health burden. Previous work has described how HCV RNA contains the RNA modification m^6^A and how this modification regulates viral infection. Yet, how this modification is targeted to HCV RNA has remained unclear due to the incompatibility of the nuclear cellular processes that drive m^6^A modification with the cytoplasmic HCV lifecycle. In this study, we present evidence for how m^6^A modification is targeted to HCV RNA in the cytoplasm by a mechanism in which WTAP recruits the m^6^A-methyltransferase METTL3 to HCV RNA. This targeting strategy for m^6^A modification of cytoplasmic RNA viruses is likely relevant for other m^6^A-modified positive-strand RNA viruses with cytoplasmic lifecycles such as enterovirus 71 and SARS-CoV-2 and provides an exciting new target for potential antiviral therapies.

## INTRODUCTION

Hepatitis C virus (HCV) is a positive-sense single-stranded RNA (ssRNA) virus that represents a significant global health burden with over 1.5 million new infections and 400,000 estimated disease related deaths annually (1). The ∼9.6 kilobase RNA genome of HCV is translated in an internal ribosome entry site (IRES)-dependent manner as a single polyprotein, which is then cleaved by host and viral proteases into ten individual viral proteins (2). These viral proteins include Core, the viral capsid protein that interacts with HCV RNA for virion production; NS5A, a key coordinator of viral RNA replication; and NS5B, the virally encoded RNA-dependent RNA-polymerase (RdRp) (2). As HCV is a positive-sense ssRNA virus, the viral RNA genome serves not only as the mRNA template for translation of the viral proteins, but also as the template for RNA replication, and as the genetic material that is packaged into virions. Thus, spatial and temporal regulation of the viral genome is essential for successful viral replication (3). Indeed, the HCV RNA genome is regulated by several RNA elements such as miRNAs, secondary structures, and RNA-binding proteins (4). In addition, we have described previously how the RNA modification N6-methyladenosine (m^6^A) plays a crucial role in regulating the HCV lifecycle (5).

The RNA modification m^6^A has now been shown to regulate infection by many viruses, through effects mediated by its presence on both viral RNA and cellular RNA (5-8). m^6^A is the most prevalent eukaryotic internal mRNA modification and regulates many aspects of RNA biology, such as mRNA stability, mRNA translation and controlling interactions with RNA binding proteins (9-12). The addition of m^6^A to mRNA, which occurs within a consensus sequence motif, DRACH (D=G/A/U, R=G/A, and H=U/C/A) is catalyzed by an enzymatic protein complex made up of the enzyme methyltransferase like 3 and its interacting cofactor methyltransferase like 14 (METTL3+14), as well as accessory proteins, such as Wilms’ tumor 1-associating protein (WTAP) which colocalize at nuclear speckles (13-17). WTAP is essential for the function, localization, and RNA targeting of the m^6^A-methyltransferase complex, and as such acts as a “central coordinator” of m^6^A (13, 16, 18). In this role, WTAP interacts with several proteins that influence targeting of the m^6^A-methyltransferase complex to specific subcellular locations and mRNAs (13, 16, 18-20). RNA modification with m^6^A can be a dynamic process with removal of m^6^A from mRNA catalyzed by enzymes such as fat mass and obesity associated protein (FTO) (21). Taken together, these m^6^A regulatory proteins have been shown to regulate diverse aspects of RNA virus infection, such as innate immune evasion, viral translation, and packaging of viral RNA into virions (5, 22-28). This regulation can also occur at the level of the host through m^6^A mediated regulation of innate immunity or viral host factors (7, 8, 29, 30). During HCV infection, we previously found that the viral RNA genome is modified by m^6^A at multiple genomic sites and is bound by the known cellular m^6^A-binding YTHDF proteins (5, 31). Further, we found that m^6^A within the coding region of the HCV E1 gene negatively regulates viral particle production by preventing the interaction of the viral Core protein with the viral RNA (5). Others have since demonstrated how m^6^A modification of HCV RNA at other sites within the genome is important for viral RNA translation by enabling recruitment of host translation factors or for promoting infection by shielding viral RNA from immune sensing by the RNA binding protein RIG-I (22, 23, 32).

The molecular mechanism of how the m^6^A-methyltransferase complex is targeted to the HCV RNA for m^6^A modification is still unclear (5). This is because the addition of m^6^A to cellular mRNA by METTL3+14 occurs in concert with RNA polymerase II-driven transcription in the nucleus, while HCV RNA replication is mediated by the RdRp NS5B and takes place in the cytoplasm (2, 3, 33, 34). Thus, m^6^A modification of HCV RNA by the m^6^A-methyltransferase complex must be occurring in a non-canonical manner in the cytoplasm. While WTAP and other members of the METTL3+14 m^6^A-methyltransferase complex are predominately localized to the nucleus, we and others have previously shown by biochemical fractionation that these proteins can be detected in the cytoplasm (5, 35). Further, METTL3 m^6^A-modification independent functions in the cytoplasm have been described (36). In fact, when the m^6^A-methyltransferase complex member ZC3H13 is depleted, biochemical fractionation and immunofluorescence microscopy revealed that multiple members of the m^6^A-methyltransferase complex, including METTL3 and WTAP, relocalize away from the nucleus (35), suggesting that these proteins may have undescribed cytoplasmic roles. Interestingly, studies of RNA viruses modified with m^6^A have demonstrated that viral infection can alter WTAP, METTL3, and METTL14 localization to the cytoplasm (28, 37, 38). Here, we investigated the hypothesis that WTAP targets METTL3+METTL14 to HCV RNA for m^6^A modification and m^6^A-mediated regulation of HCV infection. We found that WTAP is present in the cytoplasm following HCV infection and that it recruits METTL3 to HCV RNA for m^6^A modification. In addition, WTAP, like METTL3+14, negatively regulates the production of viral particles. Importantly, we also found the nuclear localization of WTAP was dispensable for m^6^A modification of HCV RNA and not required for regulation of infection. Overall, this work shows that WTAP actions in the cytoplasm control the METTL3+14-mediated m^6^A modification of HCV RNA.

## RESULTS

### HCV infection alters the subcellular localization of the m^6^A machinery accessory protein WTAP

To determine if HCV infection alters the nuclear localization of proteins in the m^6^A-methyltransferase, we fixed and stained Huh7 liver hepatoma cells that were infected with HCV or mock-infected for 48 hours and analyzed METTL3 subcellular localization by confocal microscopy. We found that METTL3 was predominantly localized to the nucleus in both mock and HCV-infected cells, with some distinct localization to the cytoplasm, and that this subcellular distribution of METTL3 did not change during infection (Fig. 1A-B). However, when we analyzed WTAP localization under the same conditions, we found that in response to HCV infection WTAP is present outside of the nucleus and that it localizes in close proximity with the HCV NS5A protein, a marker of viral RNA replication compartments (39) (Fig. 1C). In contrast, mock-infected cells show only a limited level of WTAP in the cytoplasm (Fig. 1C). Indeed, quantification of the cytoplasmic and nuclear WTAP reveals increased WTAP in the cytoplasm in HCV-infected cells (Fig. 1D). While unlike WTAP, METTL3 localization was not changed with HCV infection, the basal levels of METTL3 in the cytoplasm were higher than those of WTAP in uninfected cells (Fig. 1B-D). These data reveal that HCV infection results in increased localization of WTAP to the cytoplasm near sites of HCV replication and that METTL3 can be detected in the cytoplasm irrespective of HCV infection.

**Figure 1:**
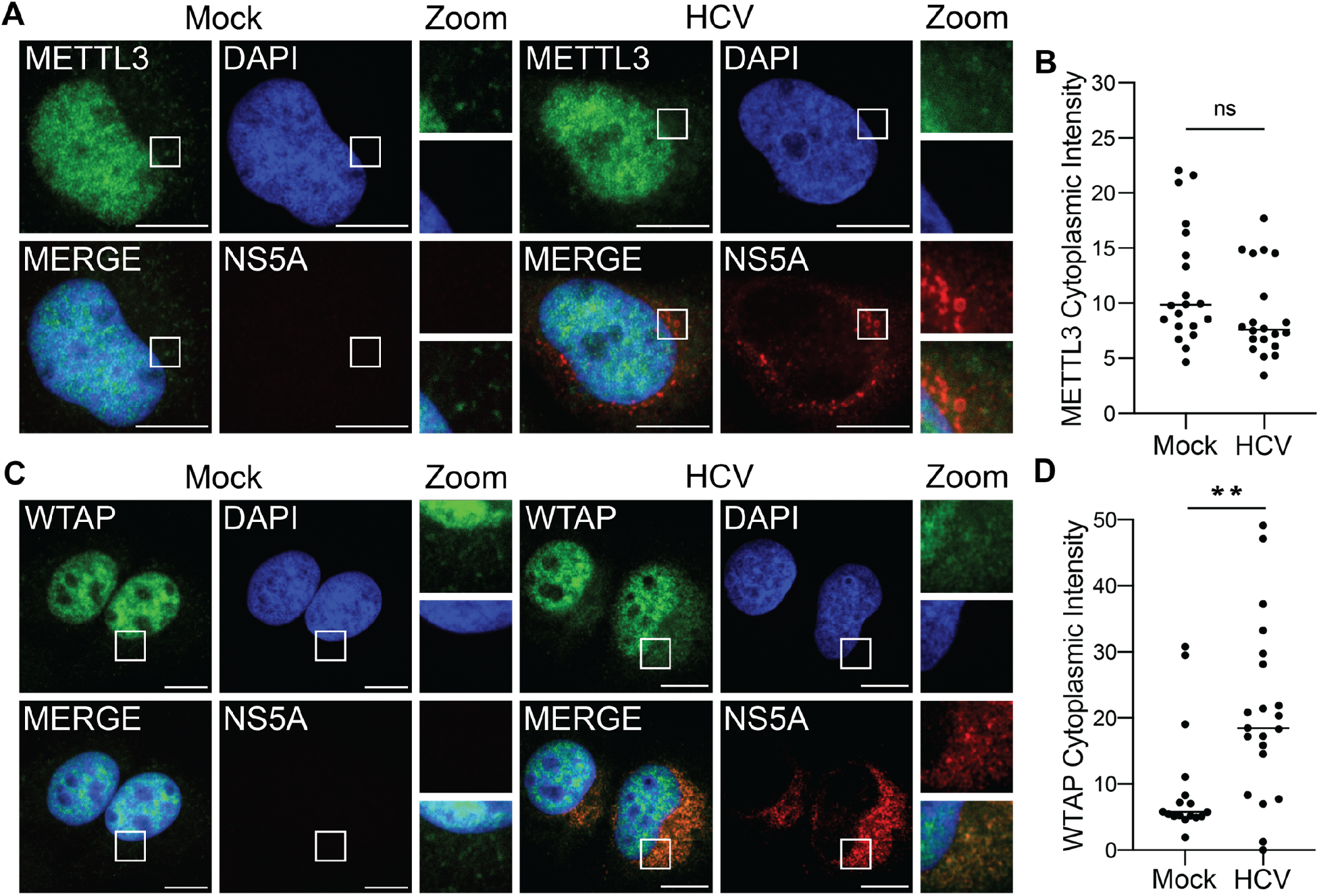
HCV infection alters the subcellular localization of the m^6^A machinery accessory protein WTAP. **(A, C)** Confocal micrographs of Mock or HCV (48 hours, MOI 0.3) infected Huh7 cells stained with DAPI and antibodies against HCV NS5A and either METTL3 **(A)** or WTAP **(C)**. Zoom is taken from area in the white box. **(B, D)** Quantification of the cytoplasmic intensity of METTL3 **(B)** or WTAP **(D)**, as described in the methods. Scale bars = 10 μM. Graph shows mean ± SD, n=21 fields. Data analyzed by Welch’s unequal variances *t*-test (* - P < 0.05, ** - P < 0.01, *** - P < 0.001, ns= not significant).

### WTAP and METTL3+14 are essential for m^6^A modification of HCV RNA

Previously we found that abrogation of m^6^A at specific sites in the E1 coding region of the HCV RNA genome, as well as depletion of METTL3+14, increases the number of infectious viral particles by promoting viral RNA interaction with Core and packaging into virions (5). As METTL3+14, as well as WTAP, are essential for m^6^A modification of cellular mRNA, we tested if they are similarly required for m^6^A modification of HCV RNA (13, 16). To accomplish this, we extracted RNA from Huh7 cells that were siRNA depleted of WTAP, METTL3+14, or non-specific control and infected with HCV for 48 hours. We then measured the m^6^A levels of previously identified m^6^A peaks or YTHDF protein binding sites on fragmented viral and host RNA by m^6^A-specific methylated RNA immunoprecipitation with qPCR (meRIP-qPCR) (Fig. 2A) (5, 7, 15). We found that depletion of both WTAP and METTL3+14 led to reduced m^6^A levels at previously characterized HCV m^6^A sites on HCV RNA (Fig. 2A-2B) (5, 23). Similarly, depletion of both WTAP and METTL3+14 led to a significant reduction in the m^6^A levels on *GOLGA3*, a transcript known to be m^6^A modified during HCV infection (Fig. 2B-2C) (7). These data demonstrate that both WTAP and METTL3+14 are essential for the m^6^A modification of HCV RNA.

**Figure 2:**
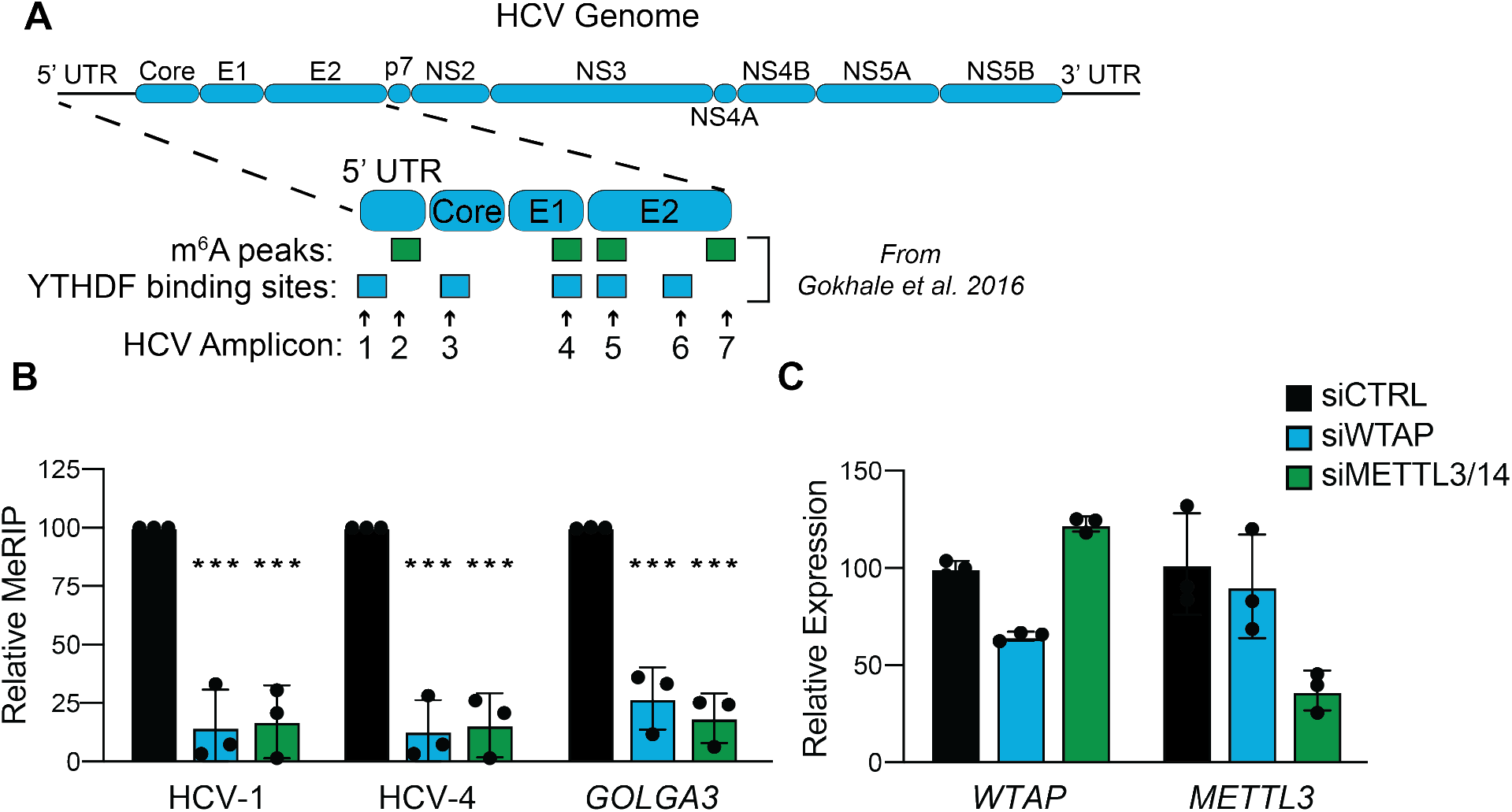
WTAP and METTL3+14 are essential for m^6^A modification of HCV RNA. **(A)** Illustration of the HCV RNA genome with amplicons measured in this study and m^6^A peaks or YTHDF protein binding sites identified in (5) are indicated. **(B)** Relative meRIP enrichment of the indicated viral or cellular amplicons from Huh7 cells treated with the indicated siRNAs and infected with HCV (48 hours, MOI 0.3), and **(C)** RT-qPCR analysis of indicated genes, relative to *18S* rRNA. For B, graph shows mean ± SD, n=3 biological replicates, while C is representative. Data analyzed by two-way ANOVA with Šidák’s multiple comparison test (* - P < 0.05, ** - P < 0.01, *** - P < 0.001, ns = not significant).

### METTL3 directly binds HCV RNA in a WTAP-dependent manner

The interaction of METTL3 with its mRNA substrates requires WTAP (16, 40). Thus, we next tested whether METTL3 directly interacts with HCV RNA and if WTAP is required for this interaction. We used ultraviolet (UV) light to cross-link protein and RNA in Huh7 cells treated with siRNA against WTAP or control and infected with HCV for 72 hours. RNA-protein complexes extracted from homogenized cells were immunoprecipitated with an antibody against METTL3, or IgG as a non-specific control, followed by capture of the bound complexes and stringent washing (Fig. 3A). RT-qPCR was then performed on extracted RNA with primers targeting previously described HCV RNA m^6^A sites or m^6^A-reader YTHDF protein binding sites (5). Immunoprecipitation of METTL3-RNA complexes enriched HCV RNA regions spanning the viral genome, while non-specific IgG control did not (Fig. 3B). Importantly, depletion of WTAP abrogated METTL3 enrichment of many of these HCV RNA regions (Fig. 3B). Together these data reveal that METTL3 directly interacts with HCV RNA and that WTAP is required for this interaction at several sites along the viral RNA genome.

**Figure 3:**
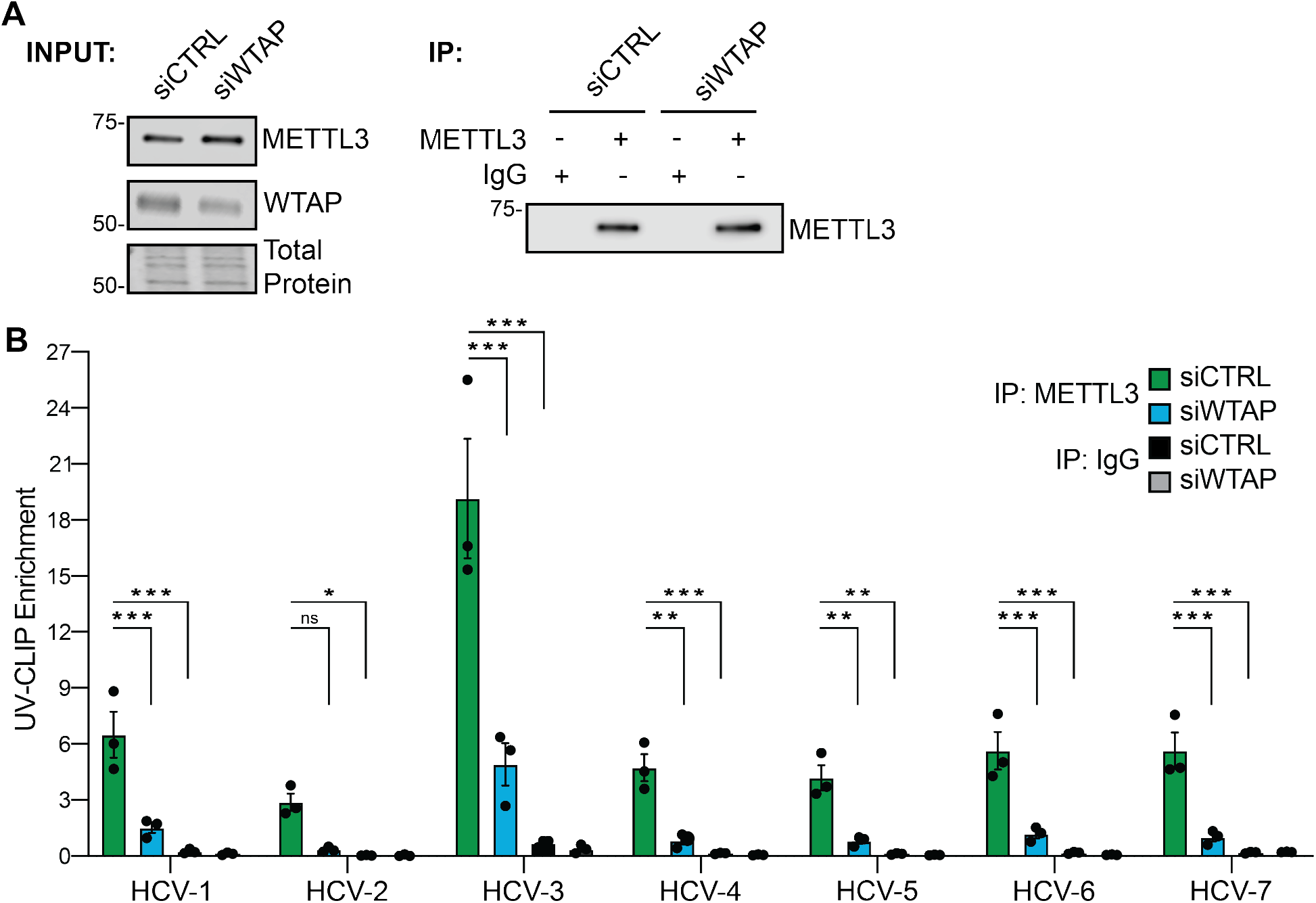
METTL3 directly binds HCV RNA in a WTAP-dependent manner. Huh7 cells were treated with the indicated siRNA and infected with HCV (72 hours, MOI 1), followed by UV-CLIP with anti-METTL3 or non-specific IgG. **(A)** Immunoblot analysis of input and immunoprecipitated UV-CLIP lysates. **(B)** Enrichment of indicated amplicons as measured by RT-qPCR. Graph shows mean ± SD, n=3 biological replicates; blot is representative of 3 independent experiments. Data analyzed by two-way ANOVA with Šidák’s multiple comparison test (* - P < 0.05, ** - P < 0.01, *** - P < 0.001, ns = not significant).

### WTAP negatively regulates HCV virion production

We previously showed that METTL3+14 negatively regulate HCV infection by decreasing the production of infectious viral particles (5). To determine whether WTAP also regulates HCV infection, we depleted WTAP by siRNA in Huh7 cells or generated Huh7 cells stably over-expressing WTAP and then used a focus-forming assay to measure the production of infectious viral particles in the cellular supernatant harvested 48 hours after HCV infection. Depletion of WTAP resulted in an increase in the production of infectious viral particles in comparison to cells treated with a non-targeting control siRNA (Fig. 4A). As we have shown before, depletion of the m^6^A-methyltransferase proteins METTL3+14 or the m^6^A demethylase FTO resulted in a similar increase, or decrease, in the production of viral particles, respectively (Fig. 4A) (5). Immunoblot analysis of cellular extracts revealed that WTAP depletion, unlike METTL3+14, resulted in decreased abundance of the HCV NS5A replicase protein as compared to siRNA control (Fig. 4B) (5). Overexpression of both WTAP and METTL3+14 reduced infectious HCV particle production relative to cells overexpressing GFP (Fig. 4C). Immunoblot analysis of lysates from infected Huh7 cells overexpressing either WTAP or METTL3+14 revealed that the abundance of the HCV NS5A protein was reduced by overexpression of these proteins, in comparison to cells overexpressing GFP (Fig. 4D).

**Figure 4:**
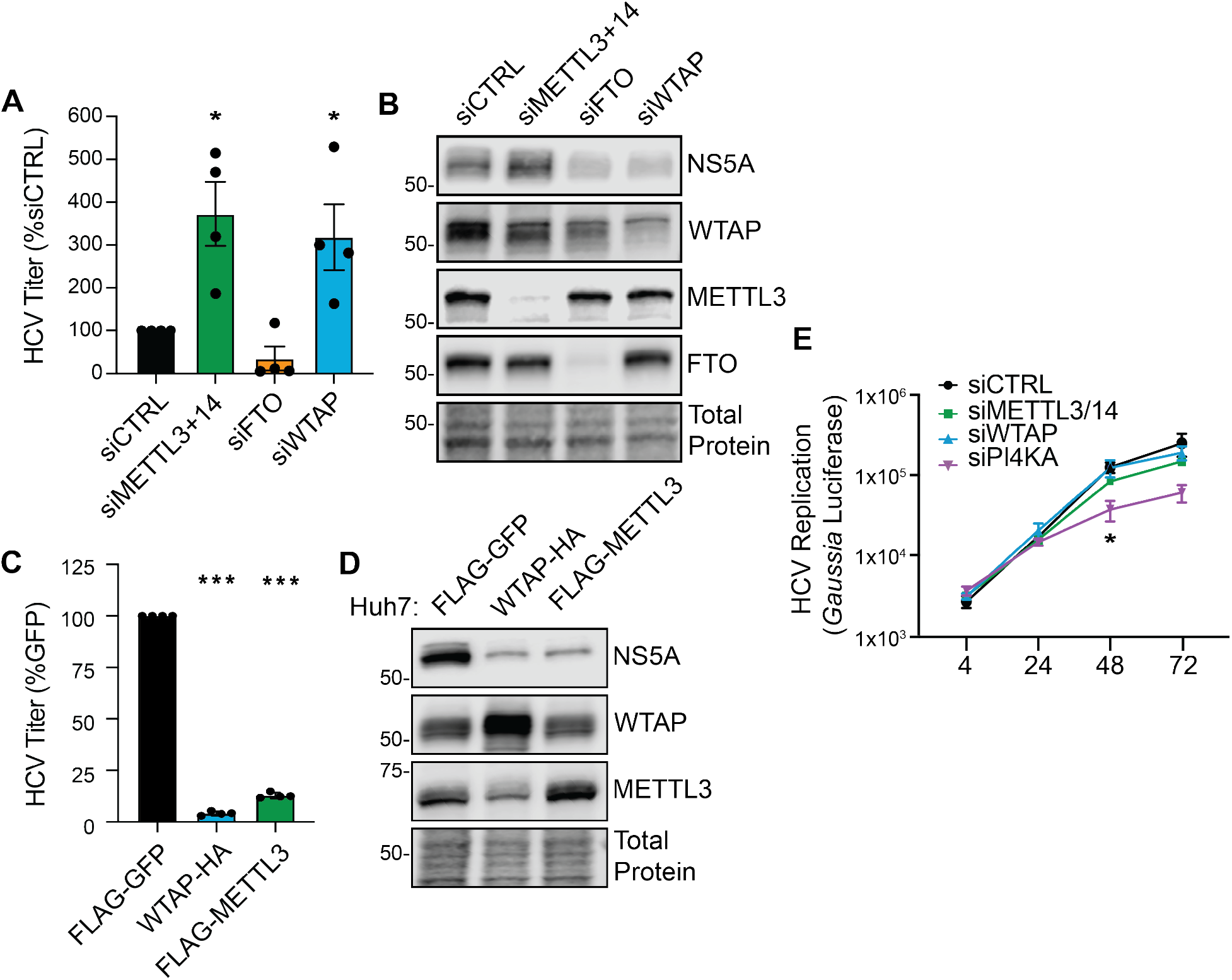
WTAP negatively regulates HCV virion production. **(A)** Focus-forming assay of supernatant harvested from HCV-infected (48 hours, MOI 0.3) Huh7 cells, as well as (**B)** immunoblot analysis of these lysates. **(C)** Focus-forming assay of supernatant harvested from HCV-infected (48 hours, MOI 0.3) Huh7 cells overexpressing the indicated proteins, as well as **(D)** immunoblot analysis of these lysates. For WTAP and METTL3, protein-specific antibodies detect both endogenous and overexpressed proteins. **(E)** *Gaussia* luciferase values from the supernatant of Huh7.5-CD81 KO cells treated with the indicated siRNA and transfected with a full-length HCV RNA containing a *Gaussia* luciferase reporter cassette, measured at indicated hour post transfection. Graphs show mean ± SD, n=3 (C, E) or 4 (A) biological replicates; blots are representative of 3 independent experiments. Data analyzed by one-way ANOVA with Šidák’s multiple comparison test (* - P < 0.05, ** - P < 0.01, *** - P < 0.001, ns = not significant).

As we previously found that METTL3+14 negatively regulate the production of infectious HCV particles, but not viral RNA replication (5), we next investigated if WTAP affected HCV RNA replication. For these experiments, we transfected HCV RNA encoding an internal *Gaussia* luciferase cassette as a reporter of viral replication (JFH1-QL/GLuc2A) into Huh7.5 cells in which the essential HCV entry factor CD81 had been deleted by CRISPR/Cas9 (Huh7.5-CD81 KO) (5, 41, 42). This allows for HCV RNA replication to be measured independent of virion production and spread. Depletion of WTAP did not alter the levels of HCV RNA replication relative to the control non-targeting siRNA over a time course (Fig. 4E). Similarly, as we have shown previously, METTL3+14 depletion did not significantly alter HCV RNA replication (5), whereas depletion of phosphatidylinositol 4-kinase alpha (PI4KA), a known host factor required for HCV RNA replication, decreased HCV RNA replication (Fig. 4E) (43). Together these data reveal that WTAP regulates the production of infectious HCV particles but does not impact viral RNA replication.

### WTAP regulation of HCV RNA m^6^A modification is independent of its nuclear localization

Cellular mRNA is m^6^A modified by the m^6^A-methyltransferase complex in the nucleus (13, 16, 17). As WTAP positively regulates HCV RNA m^6^A modification and relocalizes to the cytoplasm during HCV infection, we hypothesized that WTAP-regulation of HCV RNA m^6^A modification is independent of its nuclear localization. To test this, we generated an Huh7 cell line overexpressing WTAP lacking its described nuclear localization signal (NLS; WTAP-ΔNLS) and measured HCV m^6^A modification on fragmented viral and host RNA by meRIP-qPCR (Fig. 5A) (17). The m^6^A levels of the cellular mRNA *GOLGA3*, which we have previously shown to have increased m^6^A during HCV infection, are increased by overexpression of wild-type WTAP but not by WTAP-ΔNLS (Fig. 5A) (7). Excitingly, the m^6^A levels of multiple HCV sites across the genome are similarly increased by overexpression of either wild-type WTAP or WTAP-ΔNLS, which does not localize to the nucleus (Fig. 5A-5B). Taken together, these data reveal that WTAP regulation of HCV RNA m^6^A methylation, in contrast to cellular mRNA, can occur independent of its nuclear localization.

**Figure 5:**
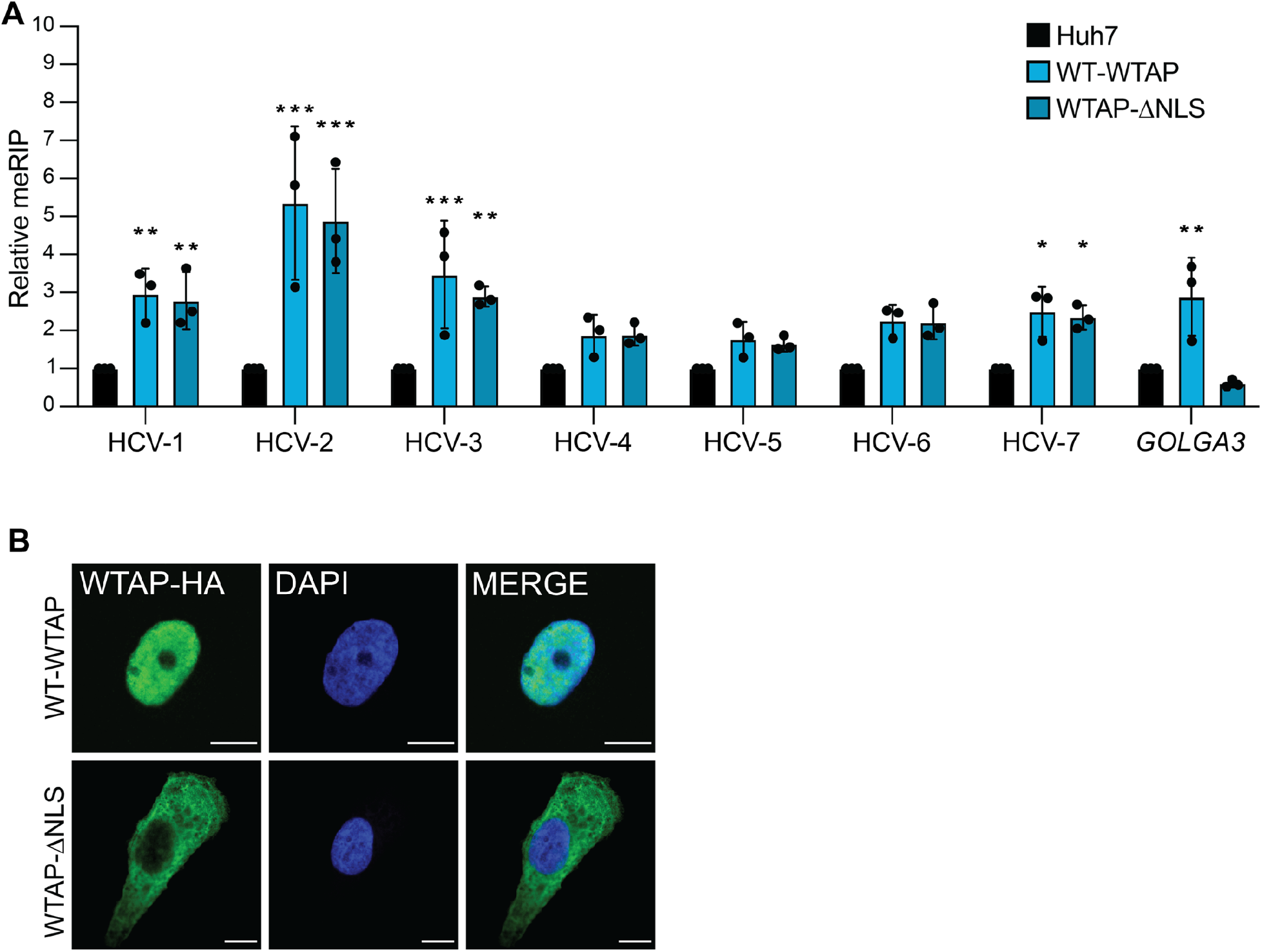
WTAP regulation of HCV RNA m^6^A modification is independent of its nuclear localization. **(A)** Relative meRIP enrichment of indicated amplicons in HCV-infected (72 hours, MOI 1) parental Huh7, wild-type-WTAP-HA, or WTAPΔNLS-HA overexpressing cells. **(B)** Confocal micrographs of Huh7 cells overexpressing WT-WTAP-HA or WTAPΔNLS-HA stained as indicated. Graph show mean ± SD, n=3 biological replicates for (A) with micrographs of localization shown in (B); Scale bars = 10 μM. Data analyzed by two-way ANOVA with Šidák’s multiple comparison test (* - P < 0.05, ** - P < 0.01, *** - P < 0.001, ns = not significant).

### WTAP regulation of HCV virion production requires METTL3 interaction but not nuclear localization

As our data reveals that WTAP regulates HCV RNA m^6^A modification and that this regulation is independent of WTAP nuclear localization, we next sought to determine features of WTAP required for regulation of HCV virion production. To accomplish this, we used Huh7 cells overexpressing wild-type WTAP, WTAP-ΔNLS, or a newly generated cell line expressing a mutant WTAP that does not interact with METTL3 (WTAP-ΔMETTL3), as seen by co-immunoprecipitation (17, 44) (Fig. 6A). We then measured the production of infectious viral particles in the cellular supernatant 48 hours after HCV infection by focus-forming assay. We found that WTAP requires its METTL3 interaction domain to negatively regulate HCV particle production (Fig. 6B). However, WTAP lacking its nuclear localization signal still reduced HCV particle production, although not as much as wild-type WTAP (Fig. 6B). Immunoblot analysis of these lysates revealed that the levels of HCV NS5A protein are decreased by both wild-type WTAP and WTAP-ΔNLS but not by WTAP-ΔMETTL3, which corroborates the results of the focus-forming assay (Fig. 6C-6D). Although WTAP is not expressed equally between the mutants, the difference in expression is not equal to the magnitude of the reduction in infectious viral particles (Fig. 6B-6E). Taken together, these data reveal that WTAP features essential for its regulation of HCV RNA m^6^A modification, but not those needed for cellular mRNA modification (Fig. 5), are required for negative regulation of HCV virion production.

**Figure 6:**
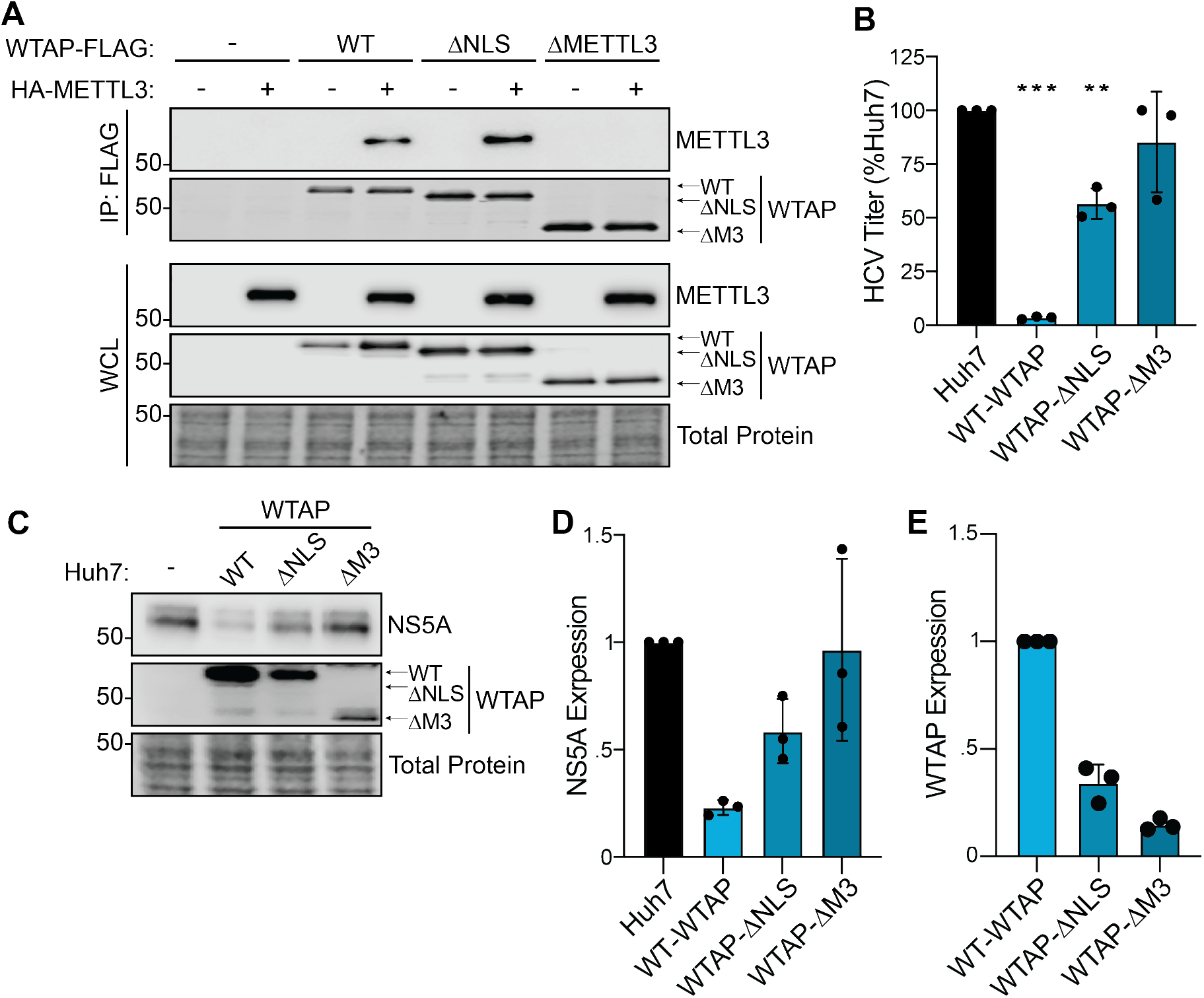
WTAP regulation of HCV virion production requires METTL3 interaction but not nuclear localization. **(A)** Immunoblot analysis of anti-FLAG immunoprecipitated lysates from Huh7 cells co-transfected with HA-METTL3 and indicated WTAP-FLAG constructs, either wild-type, ΔNLS, or ΔMETTL3 interaction. **(B)** Focus-forming assay of supernatant harvested from HCV-infected (48 hours, MOI 0.3) Huh7 cells overexpressing the indicated protein, as well as **(C)** immunoblot analysis of lysates and **(D-E)** quantification of NS5A and WTAP levels, relative total protein. Graphs show mean ± SD (n=3 biological replicates), blots are representative of 3 independent experiments. Data analyzed by one-way ANOVA with Šidák’s multiple comparison test (* - P < 0.05, ** - P < 0.01, *** - P < 0.001, ns = not significant).

## DISCUSSION

Previously, we found that the HCV RNA genome is modified by m^6^A and that both abrogation of specific m^6^A sites on the HCV RNA genome and depletion of METTL3+14 regulate viral particle production (5). Others have shown that m^6^A sites on HCV RNA also can promote viral translation or evasion of RIG-I sensing (23, 32). While these studies reveal that m^6^A on HCV RNA regulates the viral lifecycle, the mechanisms by which the HCV RNA in the cytoplasm is targeted by METTL3+14 for m^6^A modification have remained unclear. This is in part because METTL3+14 are described as functioning in the nucleus, and the mechanisms by which they are known to target mRNA for m^6^A modification would not apply to targeting HCV RNA in the cytoplasm (33, 34, 45, 46). For example, METTL3+14 localize to chromatin and interact with RNA polymerase II to m^6^A methylate mRNA in a co-transcriptional process, but HCV RNA replication does involve RNA polymerase II or occur in the nucleus near chromatin (33). While our previous biochemical fractionation experiments did reveal that METTL3+14 could be localized in the cytoplasm, and others have shown the METTL3 has cytoplasmic functions, neither total cellular nor cytoplasmic levels of METTL3+14 change with HCV infection, and so how these proteins could be repurposed for cytoplasmic addition of m^6^A was unclear (5, 36). Here, we set out to determine how METTL3+14 are targeted to HCV RNA for m^6^A modification in the cytoplasm. We found that WTAP, the METTL3+14 interacting protein that coordinates RNA targeting by the broader m^6^A-methyltransferase complex, has increased localization to the cytoplasm during HCV infection. Importantly, we found that WTAP is essential for METTL3 interaction with HCV RNA and its m^6^A modification, and that WTAP negatively regulates the production of infectious HCV particles, like METTL3+14 (5). Thus, this work reveals new insights into how the m^6^A-methyltransferase complex is repurposed to control HCV infection and highlights the contribution of WTAP in regulating HCV RNA m^6^A modification and infection.

Our work shows that proteins beyond METTL3 are important to target the m^6^A-methyltransferase complex to HCV RNA for modification. This is supported by the fact that although METTL3 preferentially modifies DRACH motifs *in vitro*, not all consensus motifs are modified in cells (14, 15, 47-50). Indeed, METTL3 was originally identified as part of a complex of proteins with m^6^A-methyltransferase activity (51, 52). This protein complex is now known to include both METTL14 and WTAP, which have specific functions in regulating m^6^A deposition (13, 16). METTL14 interacts with METTL3 and targets METTL3+14 to sites of active transcription marked by histone H3 trimethylation at lysine 36 (53). METTL3+14 then interact with WTAP (13, 16, 17), and WTAP broadly controls RNA targeting of the m^6^A-methyltransferase complex, with targeting to specific mRNAs regulated by WTAP-interacting proteins such as RBM15, VIRMA, and ZC3H13 (18-20, 35, 54). This m^6^A-methyltransferase complex then interacts with RNA polymerase II to add m^6^A to nascent mRNA (33, 34). While this mechanism for m^6^A modification drives the bulk of m^6^A on mRNA, in some cases, METTL3 can be directly recruited to mRNA transcription start sites via the protein CEBPZ (46). HCV m^6^A modification must happen differently than cellular m^6^A modification because the HCV RNA is regulated differently than cellular RNA. First, HCV transcription is mediated by the viral RNA-dependent RNA-polymerase NS5B and not RNA polymerase II and thus m^6^A modification of HCV RNA is not necessarily co-transcriptional (2). Second, HCV RNA is solely present in the cytoplasm, separate from nuclear chromatin and histones (3). As such, a unique mechanism must recruit METTL3+14 to HCV RNA. Our work shows that WTAP is important for METTL3+14 targeting to HCV RNA, that WTAP relocalizes to the cytoplasm during infection, and that m^6^A modification of HCV in the cytoplasm is driven by WTAP.

WTAP relocalization during HCV infection is likely a key factor that drives how viral RNA gets m^6^A modified in the cytoplasm; however, questions remain as to how this occurs. We do know that during infection, HCV remodels intracellular membranes to generate replication compartments (55). These compartments contain pores that are coated by nucleoporin proteins recruited from the nuclear envelope, and these nucleoporin proteins can mediate selective access for proteins involved in viral replication (56, 57). As such, we hypothesized that WTAP utilizes its NLS to access these replication compartments to facilitate m^6^A modification of HCV RNA. However, we found that the NLS of WTAP is dispensable for m^6^A modification of HCV RNA at the sites we tested (Fig. 5), while it seems to be partially required to regulate infection (Fig. 6). This suggests that WTAP regulation of HCV infection may also occur independent of its function in targeting HCV RNA for m^6^A modification. However, it is clear that WTAP recruitment to HCV RNA occurs through mechanism that does not require its NLS. It could be that a viral protein recruits WTAP to the HCV RNA. However, this would not explain how HCV infection induces WTAP localization to the cytoplasm. Interestingly, the nuclear localization of WTAP can be regulated by the cellular protein ZC3H13 (35), and a prior screen for cellular-HCV protein interactions suggested that three HCV proteins may interact with WTAP (58). Thus, it is possible that a viral protein interacts with newly translated WTAP to prevent its interaction with ZC3H13 and keep WTAP in the cytoplasm. This viral protein could then bring WTAP and METTL3+14 to HCV RNA. In support of this model, HCV E1, NS3, and NS4B, the three proteins suggested to interact with WTAP, have all been shown to broadly interact with HCV RNA and thus are candidates for bringing a WTAP/METTL3+14 complex to viral RNA for m^6^A modification (59-61). In fact, for other RNA viruses known to be m^6^A modified, such as enterovirus 71, severe acute respiratory syndrome coronavirus-2 (SARS-CoV-2), and human metapneumovirus, viral proteins do either interact or co-localize with METTL3, METTL14, or WTAP (37, 38, 62). This suggests that m^6^A-targeting may be altered during infection, indeed we have observed this previously (7). Thus, identifying which HCV and cellular proteins interact with the m^6^A-methyltransferase complex during infection will be critical to understanding how m^6^A modification is regulated during infection.

m^6^A modification of HCV RNA occurs at several positions across the genome, and several of these m^6^A sites regulate specific aspects of the HCV lifecycle (5, 23, 32). Our initial study identified sixteen high confidence m^6^A sites across the HCV RNA genome, and three of these sites have unique functions during HCV infection (5, 23, 32). Our understanding of how m^6^A modification at each of these sites occurs in relation to each other is limited. It may be that there are unique mechanisms that recruit METTL3+14 to viral RNA to m^6^A modify specific sites, with differing effects on the HCV lifecycle. This possibility may explain why METTL3+14 and WTAP depletion differentially affect NS5A expression, although it is unclear how m^6^A could both positively regulates IRES-mediated translation and negatively regulate the levels of HCV proteins (Fig. 4) (23). Importantly, current methods to identify m^6^A do not allow for specific mapping of the m^6^A profile for each copy of the viral RNA during infection. This m^6^A profile on individual viral RNAs may regulate distinct viral processes, such as translation, transcription, or virion production. As such, it may be that the HCV RNA molecules involved in active translation may have one set of m^6^A sites modified, whereas those involved in viral packaging have a different set of m^6^A sites modified. Each of these m^6^A profiles could arise from different viral RNA targeting factors. In fact, WTAP enhancement of m^6^A modification does not appear to be uniform for all tested regions of HCV RNA (Fig. 5). Thus, additional viral RNA targeting factors may be required for modification of particular m^6^A sites.

Overall, this study reveals that WTAP is an important regulator of m^6^A modification of a cytoplasmically localized RNA. Specifically, WTAP regulates HCV RNA m^6^A modification and as such it regulates virion production. Importantly, this regulation by WTAP is independent of its ability to localize to the nucleus. Thus, this work supports of model by which HCV infection induces WTAP localization changes to mediate cytoplasmic m^6^A modification of viral RNA. Studies of how methylation of specific HCV m^6^A sites are controlled and how HCV RNA m^6^A modification is regulated throughout the viral lifecycle will undoubtedly provide insight into the mechanisms involved. Our work reveals that we still have much to learn of the processes that govern m^6^A methylation, an RNA regulatory mechanism critical in cellular differentiation, numerous cancers, and infection by an ever-growing list of viruses, including those of global health concern such as SARS-CoV-2 and members of the *Flaviviridae*.

## ACKNOWLEDGEMENTS

We thank those colleagues who generously provided reagents including Dr. Matthew Evans, Dr. Michael Gale Jr., Dr. Stanley Lemon, and Dr. Charlie Rice; the Duke Functional Genomics Core, the Duke Light Microscopy Core Facility, and members of the Horner Lab for valuable feedback and discussion. This work was supported by Burroughs Wellcome Fund (S.M.H, M.T.S.) and National Institutes of Health grants R01AI125416 (S.M.H, M.T.S., K.M.B) and T32CA00911 (M.T.S, K.M.B).

## METHODS

### Cell Culture

Huh7, Huh7.5 (gift of Dr. Michael Gale Jr., University of Washington (63)), Huh7.5 CD81 KO (gift of Dr. Matthew Evans, Icahn School of Medicine at Mount Sinai (64)) and 293T cells were grown in Dulbecco’s modification of Eagle’s medium (DMEM; Mediatech) supplemented with 10% fetal bovine serum (HyClone), 25 mM N-2-hydroxyethylpiperazine-N’-2-ethanesulfonic acid (Thermo Fisher), and 1X non-essential amino acids (Thermo Fisher), referred to as complete DMEM (cDMEM). Cells were verified using the Promega GenePrint STR kit (DNA Analysis Facility, Duke University) and as mycoplasma free by the LookOut Mycoplasma PCR detection kit (Sigma-Aldrich).

### Plasmids

The following plasmids were generated by subcloning polymerase chain-reaction (PCR) generated amplicons from the indicated oligonucleotides from Table 1 into pEFtak or pLEX vector using In-fusion recombinase (Takara) according to manufacturer’s instructions: pEFtak-WTAP-FLAG, pEFtak-WTAPΔNLS-FLAG, pEFtak-WTAPΔMETTL3-FLAG, pEFtak-HA-METTL3, pLEX-WTAP-HA, pLEX-FLAG-METTL3, pLEX-WTAPΔNLS-HA, pLEX-FLAG-GFP and pLEX-WTAPΔMETTL3-HA. pJFH1-QL/GLuc2A was a gift of Dr. Stanley Lemon (University of North Carolina at Chapel Hill (42)).

**Table 1.**
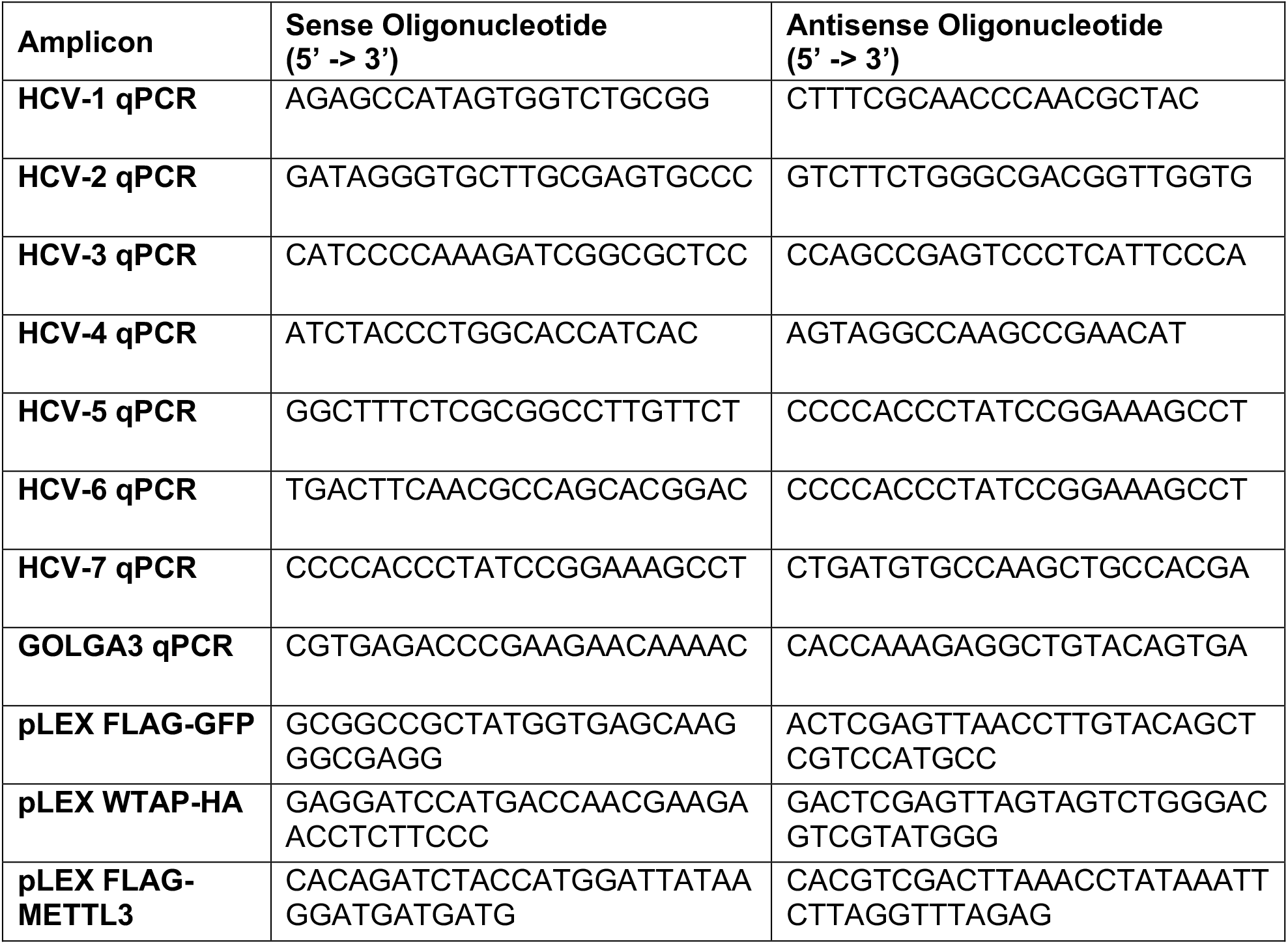
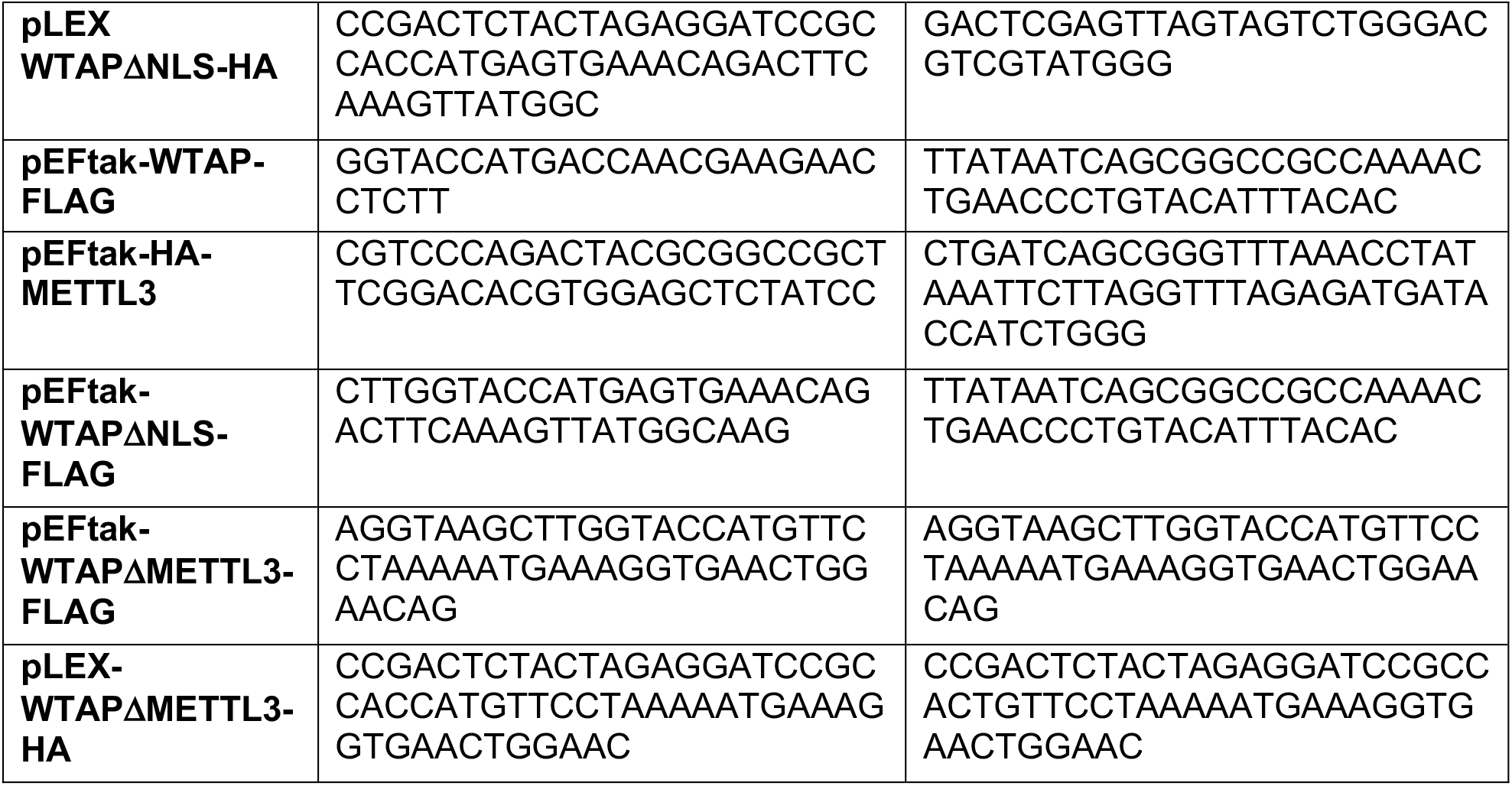

### *In Vitro* Transcription

Generation of HCV luciferase reporter RNA was accomplished with the MEGAscript T7 transcription kit (Invitrogen) using XbaI-linearized and Mung Bean Nuclease-treated (enzymes from New England Biolabs) JFH1-QL/GLuc2A plasmid (42), following the manufacturer’s instructions.

### Viruses

Infectious stocks of a cell culture-adapted strain of genotype 2A JFH-1 HCV (JFH-1 M9 (65)) were generated in Huh7.5 cells. Measurement of viral titers and virion production from infected supernatants was performed in Huh7.5 cells by focus-forming assay (FFA), as previously described (65). For viral infections, cells were incubated in a low volume of serum free DMEM containing virus at the indicated multiplicity of infection (MOI) for 3 hours, following which cDMEM was replenished.

### Cell Line Generation

293T cells were transfected with pLEX-WTAP-HA, pLEX-FLAG-METTL3, pLEX-WTAPΔNLS-HA, or pLEX-WTAPΔMETTL3-HA and the viral packaging plasmids psPAX2 and pMD2.G (Addgene #12260 and #12259; gift of Duke Functional Genomics Facility) and supernatant was harvested and filtered with an 0.22 μm filter 72 hours after transfection. This filtered supernatant was then used to transduce Huh7 cells for 24 hours. Transduced cells were then placed in 2 μg/μL puromycin for 72 hours and then overexpression validated by immunoblotting, as described below. After selection and validation, cell lines were maintained in 1 μg/μL puromycin cDMEM until the time of experimentation.

### siRNA Treatment

Cells were transfected with siRNA against indicated targets using Lipofectamine RNAiMAX (Thermo Fisher) according to the manufacturer’s protocol for 24 hours prior to experimental infection or treatment. Depletion of siRNA targets was confirmed by immunoblot analysis. siRNAs (Qiagen) used included siWTAP (SI00069853), siMETTL3 (SI04317096), siMETTL14 (SI00459942), siFTO (SI04177530), siPI4KA (SI02777390) and siCTRL (1027281).

### Luciferase Assay

Huh7.5 CD81 KO cells seeded into 12 well plates were transfected with siRNAs as described above. Cells were then transfected with 1 μg of JFH1-QL/GLuc2A *in vitro* transcribed RNA (1 μg) using polyethylenimine (Polysciences, Inc.). Supernatants were harvested and mixed with Renilla luciferase buffer and substrate, and luciferase values were measured according to manufacturer’s instructions (Renilla Luciferase Assay System, Promega) using a BioTek Synergy 2 microplate reader.

### Immunoblotting

Cells were lysed in a modified radio immunoprecipitation assay (RIPA) buffer (10 mM Tris pH 7.5, 150 mM NaCl, 0.5% sodium deoxycholate, 1% Triton X-100) supplemented with protease inhibitor cocktail (Sigma-Aldrich) and phosphatase inhibitor cocktail (Millipore), and supernatants were collected after centrifugal clarification. Quantified protein, as determined by Bradford assay (BioRad), was resolved by SDS-PAGE, transferred to nitrocellulose or polyvinylidene difluoride (PVDF) membranes using the Turbo-transfer system (BioRad), stained with REVERT total protein stain (Licor Biosciences) and blocked with 3% bovine serum albumin (BSA) (Sigma-Aldrich) in phosphate buffered saline (PBS) with 0.1% Tween (PBS-T). Membranes were probed with specific antibodies, washed with PBS-T and incubated with species-specific HRP conjugated antibodies (Jackson ImmunoResearch), washed again with PBS-T, and treated with Pico PLUS enhanced chemiluminescent (ECL) reagent (Thermo Fisher). The signal was then captured by using a LICOR Odyssey FC.

### Protein Immunoprecipitation

50-100 μg of protein extracted as above was incubated with 25 μL anti-FLAG M2 magnetic beads (Sigma-Aldrich) in modified 1X RIPA in a total volume of 300 μL at 4°C overnight with rotation. Beads were washed 3 times in PBS and eluted in 40 μL 2X Laemmli Buffer with 1:20 β-mercapto-ethanol (BioRad) at 95°C for 5 minutes. Eluates were resolved by SDS-PAGE and immunoblotting, as described above.

### RT-qPCR

Total cellular RNA was extracted using the Qiagen RNeasy kit (Life Technologies) or TRIzol extraction (Thermo Fisher). RNA was then reverse transcribed using the iSCRIPT cDNA synthesis kit (BioRad) as per the manufacturer’s instructions. The resulting cDNA was diluted 1:5 in nuclease-free distilled H2O. RT-qPCR was performed in triplicate using the Power SYBR Green PCR master mix (Thermo Fisher) and the Applied Biosystems QuantStudio 6 Flex RT-PCR system. Primer sequences for RT-qPCR are listed in Table 1.

### MeRIP

For meRIP, total RNA was extracted from cells using TRIzol (Thermo Fisher) according to the manufacturer’s protocol and diluted to equivalent concentrations. Then, meRIP was performed as previously described (8). Following meRIP, cDNA from the input and immunoprecipitated RNA fractions was generated and analyzed by RT-qPCR as described above. Relative m^6^A level for each transcript was calculated as the percent of input in each condition normalized to that of the respective positive control m^6^A RNA spike-in, as described (8). Percent change of enrichment was calculated with siCTRL samples normalized to 100.

### UV-CLIP

UV-CLIP was adapted as a modified version of formaldehyde CLIP (40). Briefly, Huh7 cells were plated in 10 cm dishes, treated with siRNA, and HCV-infected as described above. For UV crosslinking, supernatant was removed and replaced with 2.5 mL of 4°C PBS. Plates were then irradiated with 150 mJ/cm^2^ 254 nm UV and then cross-linked cells were harvested in 500 μL CLIP-RIPA buffer (50 mM Tris-HCL pH 7.4, 100 mM NaCl, 1% CA-630, 0.1% SDS, 0.5% sodium deoxycholate) supplemented with 1 mM dithiothreitol (DTT), protease inhibitor, and RNAseIN+ (Promega). Next, the cross-linked cells were passed through a Qiashredder column (Qiagen) twice to generate homogenized lysates. These lysates were incubated with 2 μL Turbo DNAse I and 1 μL 1:2000 diluted RNAse I for fragmentation for 25 minutes at 37°C with constant agitation and then clarified by centrifugation (enzymes from New England Biolabs). Equivalent amounts of lysates were then precleared for 4 hours using protein A beads (Thermo Fisher) that were preblocked (1 μg of yeast tRNA and 1% BSA per 100 μL of beads in a total volume of 750 μL CLIP-RIPA buffer). RNA and protein inputs were reserved from these lysates and prepared as follows: for the RNA input, crosslinks were removed by incubating equal amounts (50 μg) of precleared lysate in a total volume of 250 μL of CLIP-elution buffer (50 mM Tris-HCL pH 7.4, 5 mM EDTA, 10 mM DTT, 1% SDS, 1% RNAseIN+, 1:100 Proteinase K) and incubated at 50°C for 1 hour with constant agitation, with RNA extracted using TRIzol-LS (Thermo Fisher) and reserved for RT-qPCR; for the protein input, ∼10 μg was reserved for immunoblotting. The remaining precleared lysates were divided and incubated at 4°C for >12 hours with either METTL3-bound or IgG-bound pre-blocked protein A beads in 1 mL of CLIP-RIPA + 2 μL RNAseIN+. Then, these samples were washed 5X (CLIP-RIPA +1 M NaCL & 1 M Urea) and 1X (CLIP-RIPA) followed by resuspension in 100 μL CLIP-elution buffer without Proteinase K. A portion of this eluate (10 μL) was reserved for immunoblotting, while CLIP-elution buffer (160 μL of CLIP-elution buffer) + Proteinase K (2 μl) was added to the remaining 90 μL of beads, which were incubated for 1 hour at 50°C with constant agitation, and RNA was extracted using TRIzol-LS and this and the input were analyzed by RT-qPCR.

### Immunofluorescent Microscopy

Cells were fixed in 4% paraformaldehyde in PBS, permeabilized with 0.2% Triton X-100 in PBS, and blocked with 10% FBS in PBS. Slides were stained with indicated antibodies, and incubated with conjugated Alexa Fluor secondary antibodies (Life Technologies) and mounted with ProLong Diamond + 4’,6-diamidino-2-phenylindole (Invitrogen). Imaging was performed on a Zeiss 880 laser scanning confocal microscope, using a 63x/1.25 oil objective using 405, 488, 561 and laser lines at a 4x optical zoom with pinholes set to 1 AU for 561 (Light Microscopy Core Facility, Duke University), or a Leica DM4B widefield fluorescent microscope. Gain and offset settings were optimized, and final images were taken with line averaging of 4. All images were processed with NIH Fiji/ImageJ (66). To quantify the Cytoplasmic:Nuclear ratios of proteins, 7 fields from each biological replicate with at least 5 cells each (21 fields total, >100 cells per condition) were analyzed in NIH Fiji/ImageJ (66) using the Intensity Ratio Nuclei Cytoplasm Tool, RRID:SCR_018573, with protein (METTL3 or WTAP) signal intensity demarcated by the tool, and then calculated as an average of all cells in each field.

### Antibodies

Antibodies used in this study and their applications include HCV NS5A 9e10 (gift of Dr. Charles Rice; Immunoblot, FFA, Immunofluorescence), FLAG-HRP (Sigma-Aldrich, A8592; Immunoblot), HA (Sigma-Aldrich H6908; Immunoblot), METTL3 (Novus Biologicals, AB_2687437; Immunoblot), METTL3 (Abcam, ab195352; Immunofluorescence, Immunoprecipitation) FTO (Abcam, ab92821; Immunoblot), WTAP (Proteintech, AB_10859484; Immunoblot), WTAP (Abcam, ab195380; Immunofluorescence), and non-specific rabbit IgG (Cell Signaling Technologies, 2729S; Immunoprecipitation).

### Statistical Analysis

Statistical analysis was performed using Graphpad Prism 9. Data appropriate statistical test were performed including 1- and 2-way ANOVA with post-hoc testing (Figures 2, 3, 4, 5, 6) or Welch’s t-test (Figure 1). Values are presented as mean ± standard deviation of the mean for biological replicates (n=3, or as indicated). * - P < 0.05, ** - P < 0.01, *** - P < 0.001.

